# StereoGene: Rapid Estimation of Genomewide Correlation of Continuous or Interval Feature Data

**DOI:** 10.1101/059584

**Authors:** Elena D. Stavrovskaya, Tejasvi Niranjan, Elana J. Fertig, Sarah J. Wheelan, Alexander Favorov, Andrey Mironov

## Abstract

**Motivation:** Genomics features with similar genomewide distributions are generally hypothesized to be functionally related, for example, co-localization of histones and transcription start sites indicate chromatin regulation of transcription factor activity. Therefore, statistical algorithms to perform spatial, genomewide correlation among genomic features are required.

**Results:** Here, we propose a method, StereoGene, that rapidly estimates genomewide correlation among pairs of genomic features. These features may represent high throughput data mapped to reference genome or sets of genomic annotations in that reference genome. StereoGene enables correlation of continuous data directly, avoiding the data binarization and subsequent data loss. Correlations are computed among neighboring genomic positions using kernel correlation. Representing the correlation as a function of the genome position, StereoGene outputs the local correlation track as part of the analysis. StereoGene also accounts for confounders such as input DNA by partial correlation. We apply our method to numerous comparisons of ChIP-Seq datasets from the Human Epigenome Atlas and FANTOM CAGE to demonstrate its wide applicability. We observe the changes in the correlation between epigenomic features across developmental trajectories of several tissue types consistent with known biology, and find a novel spatial correlation of CAGE clusters with donor splice sites and with poly(A) sites. These analyses provide examples for the broad applicability of StereoGene for regulatory genomics.

**Availability:** The *StereoGene* C++ source code, program documentation, Galaxy integration scripts and examples are available from the project homepage http://stereogene.bioinf.fbb.msu.ru/

**Supplementary information:** Supplementary data are available online.

## Introduction

Modern high throughput genomic methods generate large amounts of data, which can come from experimental designs that compare tissue-specific or developmental stage-specific phenomena.

An important challenge of genome-wide data analysis is to reveal and assess the interactions between biological processes, *e.g.* chromatin profiles and gene expression. An emerging approach to this challenge is to represent the biological data as functions of genomic positions (we use terms *profile* or *track* for the functions) and to estimate correlations between these functions.

In recent years, the bioinformatics community has actively developed methods for assessment of colocalization of genomic features [6, 9, 14, 28, 31]. The features are typically represented as a set of intervals on the genome (genes, repeats, CpG islands, etc.), as point profiles (binding sites, TSS, splice sites), or as continuous (numeric) profiles (coverage of expression, ChIP, etc from high-throughput sequencing experiments). A common approach to investigating genomic features is to represent these features as intervals, computed from the original continuous coverage data using a threshold or more sophisticated algorithms [32]. The tracks resulting from this discretization are sensitive to algorithm parameters, including thresholds, and therefore are unable to account different levels of genomic coverage or gene usage.

In addition, most genome-wide correlation algorithms [*e.g.*, 14, 31] compare genomic features at identical genomic coordinates (called overlapping coordinates). However, biologically regulatory relationships may often occur between features within a neighborhood of genomic coordinates (called adjacent coordinates). For example, gene expression profiles (RNA-seq coverage) correlate with transcription factor binding sites or chromatin state in nearby promoter regions or distant enhancer regions. Interval and point-based approaches developed for genome-wide correlation account for associations between adjacent coordinates by estimating distance-based statistics [6, 9, 14].

Here, we propose a fast universal method – *StereoGene* – to correlate numeric genomic profiles. The data can be genomewide tracks with discrete features (e.g. intervals) or continuous profiles, e.g. coverage data. The method is based on kernel correlation (KC), which provides an estimate of spatially smoothed correlation of two features. The statistical significance of correlations with *StereoGene* are evaluated by a permutation-based test. *StereoGene* provides additional functionality, including a track representing correlation as a function of genomic coordinate (called the local correlation); calculation of positional cross-correlation function; account for genomewide confounders by partial correlation. Our implementation is computationally efficient: the calculation of the KC with permutations for a pair of profiles over the human genome takes approximately 1-3 minutes on a personal computer. We demonstrate the effectiveness of *StereoGene* for estimation of genome-wide epigenetic profile data correlations pairwise correlations between all human samples in the Roadmap Epigenomics Project [3] dataset and on other open data. These examples describe some potential applications of *StereoGene* for regulatory genomics to provide a template for its broad utility.

## Materials and Methods

### Kernel correlation

We consider each genomic feature as a numeric function (profile) of the genomic position *x*. The standard Pearson correlation of two profiles *f* = *f* (*x*) and *g* = *g* (*x*) is defined as:

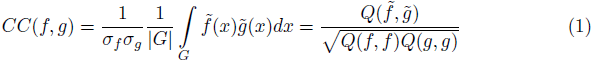

where 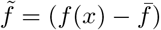, 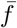 is the mean value of *f*; *σ_f_* is the standard deviation of *f*, *Q*(*f, g*) = ∫*_G_ f* (*x*)*g*(*x*)*dx*; the integration is performed over the genome *G*. The Pearson correlation relates profile values on exactly the same genomic positions. In biological systems, the relationships of values at proximal but non-overlapping (in genomic coordinates) positions are also important. These correlations may be result from transcriptional regulation, chromatin looping, or other interactions. To account for correlations between profiles at proximal coordinates, we generalize the Pearson correlation from eq. 1 to the covariation integral as follows:

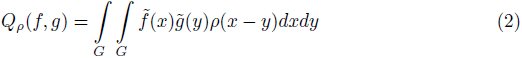

where *ρ*(*x - y*) reflects the common sense expectations of the interaction of features at adjacent positions. Formally, it is a function of the distance *x - y* between the interacting positions. In the case *ρ*(*x - y*) = *δ*(*x - y*), we get the standard Pearson correlation integral as in eq. 1. In theory, any non-negative kernel function can be used. The default kernel we use is the Gaussian 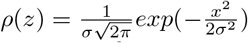, it is the most intuitive representation of the closer is the position, the more important it is. The *σ* of the Gaussian reflects the interaction scale and it a user-defined parameter with a reasonable default of 1000bp.

Based on the *Q* covariation value (eq. 2), we introduce the kernel correlation *KC* defined as:

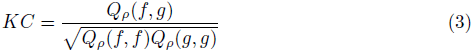

The two-dimensional integral *Q_ρ_*(*f, g*) can be rapidly calculated using a complex Fourier transform (See Supplementary File 1, Section 1).

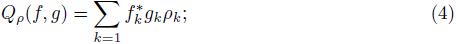

where 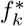*, gk, ρk* are Fourier coefficients; *^∗^* means complex conjugate. The value *KC*(*f, g*) satisfies the inequality: *-*1 *≤ KC* (*f, g*) *≤* 1. The marginal values 1 and *-*1 correspond to *f* = *g* and *f* = *-g* (see Supplementary File 2, Section 7.3 for the test) Fourier transform can be calculated by the discrete Fast Fourier Transform (FFT) algorithm [17], and therefore has computational cost of *O*(*|G| ·* log *|G|*) where *|G|* is the length of the genome.

### Cross-correlation

Sometimes, in addition to the overall value of the correlation, an investigator needs information about its local structure, e.g. either the value emerges from a strong position-to-position overlap of it comes from a smooth interaction of neighboring positions and what is the scale of interaction if it exists. To address the questions for our two profiles, *f* (*x*) and *g*(*x*), *StereoGene* calculates the cross-correlation function *c*(*x*) as follows.

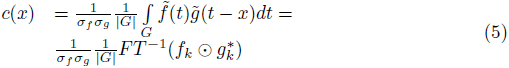

where *F T ^-^*^1^ means the inverse Fourier transform and ⊙ is elementwise product of *fk* and 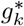 vectors of the Fourier coefficients.

### Local correlation profile

The correlation itself shows the similarity of the features at the scale of the genome. The cross-correlation function (see above) reflects the fine-scale structure of correlation. The distribution of the correlation as a function of the genomic position is also relevant to determine the nature of interactions. To provide this information, *StereoGene* generates a new track that describes the local kernel correlation of two original profiles as a function of the genomic position, called the “local correlation (LC)”.

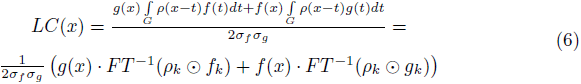

Note that the value of *LC* is not restricted by *±* 1 boundaries and can take any values. The scale of *LC* depends on the data nature, the direct comparison of *LC* values makes sense only inside one *LC* track. To give the user ability to select regions with significant enrichment of *LC*, *StereoGene* outputs the FDR *LC* value (see Statistical significance). Standard peak calling tools [e.g. MACS, 32] can be applied to the *LC*. The result is suitable for gene set enrichment analysis.

### Partial correlation

Nonrandom correlation of the two profiles may occur due to their correlation with a third profile (confounder) that systematically biases both signals (e.g. level of mapability). An example of such confounding would be the case with ChIP-seq for a sample with the signal from two antibodies (the profiles to correlate) and a common input track (the confounder). *StereoGene* can computationally exclude such a confounding using the partial correlation (projection) approach. For this calculation, *StereoGene* correlates the projections both profiles in the subspace, that is orthogonal to the profile *a* of the confounder as follows:

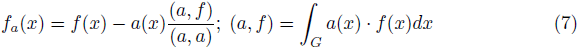

where (*a, f)* means a scalar product of the functions *a, f*. Then the *KC*, the Local correlation track, and the Cross-correlation between two projection is calculated in a standard way. The statistical significance (see below) is calculated as for a regular two-way comparison.

### Statistical significance

All the calculations we described above are executed independently in large (we recommend a size of 100kb..1M) windows (see Fig. 1). This approach allows the FFT transform to be really fast and at the same time, it provides us with the statistical significance of all the observations. We apply a permutation test to obtain significance for the computed correlation coefficients. Specifically, the correlations (foreground distribution) are calculated in a set of pairs of windows with the same genome positions on the tracks we compare (matched windows). To obtain the null distributions of the values, a shuffling procedure is used that randomly matches windows on one profile to the windows on another profile and then the correlations (background distribution) are computed for these randomly matched window pairs in the same way as they are calculated for the original (matched) window pairs. The statistical significance for *KC* is provided by a Mann-Whitney test of these two sets of values. The *F DR* for local correlation is estimated by using the background distribution as null-hypothesis and the foreground as the signal.

**Figure 1.**
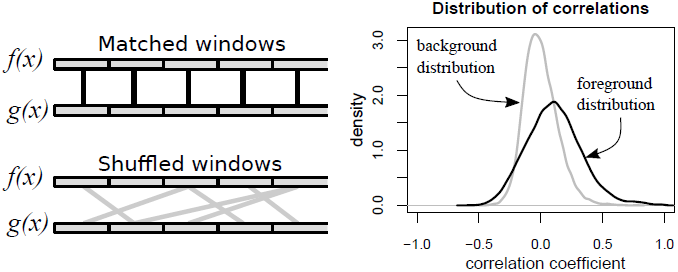
The procedure that is used for the estimation of p-values for kernel correlation and for the estimation of FDR values for the local correlation is based on shuffling of windows. Left pane: Shuffling procedure. Right pane: Background and foreground distributions.

### Program implementation

*StereoGene* is implemented as a command-line tool, and it is distributed as C++ source code under MIT 2.0 license. *StereoGene* processes the input data in two passes. On the first pass, *StereoGene* converts input profiles to an internal binary format and saves the binary profiles for the future runs. The second pass does the Fourier transforms as well as permutations and calculates all the correlations and statistics. If a project refers to a track that has its binary profile already calculated and the parameters have not been changed, *StereoGene* omits the first pass and reuses the saved profiles. The time required for the first pass depends on the input file size. On a standard computer, for a typical ChIP-Seq track, the first pass takes from a few seconds up to 1-2 minutes. The second pass takes less than half a minute on the human genome.

#### Input

As input, *StereoGene* accepts two or more input files in one of the standard genomic tracks formats: BED, WIG, BedGraph, and BroadPeak. If more that two track files are provided, *StereoGene* makes all the pairwise comparisons. *StereoGene* can take a linear model, which combines a number of profiles to get one of the tracks to compare, as an input. For a batch processing *StereoGene* accepts a text file containing a list of the tracks. A linear combination of input tracks (model) described by a text file can be used instead of a track.

#### Output

*StereoGene* reports the kernel correlation (*KC*) over all the genome; *KC* values for matched and for shuffled windows, averaged kernel correlation over the matched windows, and p-value. *StereoGene* produces the following files: the foreground and background distributions for *KC*; the cross-correlation function; the local correlation track; table of FDR values for the local correlation values; and some additional files. The information can be presented on the whole genome as well as by chromosomes.

#### *StereoGene* command-line run

The only parameter that does not have a default value and thus is required for a run is a file with the chromosome names and lengths. For the partial correlation, the confounder track should be defined. All other parameters are optional and have reasonable defaults. The less technical of them are the window size and the kernel width. Detailed information about input and output files and parameters is presented in the program documentation at the *StereoGene* homepage http://stereogene.bioinf.fbb.msu.ru/. The homepage also contains an archive with the example run scripts along with all the necessary files. A general program description is presented in the Supplementary File 2.

#### Visualization and interface

For a quick and intuitive depiction of results, the *StereoGene* provide an optional mode that prepares an R script, which represents the output in a multipanel plot (for an example, see Supplementary File 1, Section 2). The first panel displays foreground and background distributions of genomic windows by the kernel correlation. If the foreground distribution is shifted to the right of the background distribution, the plot represents the positive correlation, and the left shift shows anti-correlation. The significance of the observation is represented by the Mahn-Whitney test that is described in the Statistical Significance subsection. The second panel, which is the cross-correlation function, represents spatial relationships between the tracks. The third panel represents the local correlation distribution for the observed (foreground) and the null (background) local correlation distributions and the FDR q-values. We provide two tool definition files to use *StereoGene* in Galaxy [1], one to compute and to visualize the correlation of a pair of tracks, and another to compute and to visualize the partial correlation given a confounding track.

### Gene set analysis by the local correlation track

To detect the gene sets that are overrepresented around the areas of high local correlation of a pair of tracks, we do the following. We take the local correlation track (*.wig *StereoGene* output file) convert the track to the BED format using bedops software, version 2.4.16 [21] and selected 3000 of the highest peaks using MACS-1.4.2 [32] with the default parameters. Next, we select the genes whose transcription start sites fell within 5kb of the correlation peak. The resulting list of genes is mined for biological enrichment using DAVID 6.7 software [12]. Eventually, we obtain a list of gene-related terms (gene sets), that are significantly overrepresented around the high local correlation regions, with some statistical measures (adjusted p-value, FDR) for each term.

### Data source

Data from the Roadmap Epigenomics Project [3] are downloaded from the Human Epigenome Atlas (http://www.genboree.org/). Data for FANTOM4 CAGE clusters [24] are obtained from the UCSC website (RIKEN CAGE tracks, GEO accession IDs are GSM849326 for nucleus GSM849356 for cytosol in H1 Human Embryonic Stem Cell Line, RRID:CVCL 9771). The datasets with the tracks are listed in Supplementary File 1, Section 10.

## Results

*StereoGene* enables a variety of genome-wide correlation techniques to account for different types of interrelationships between pairs of continuous genomic features. We summarize the major types of correlations enabled by *StereoGene* in Table ??.

## *StereoGene* application examples

### Human Epigenome Atlas Pairwise Correlation Anthology

To demonstrate the wide-range of applications of *StereoGene*, we have built a pipeline that applies *StereoGene* to the Human Epigenome Atlas in the Roadmap Epigenomics Project [3]. This database is comprehensive, containing 2423 different data types for 186 different tissues (263077 pairs total). For our analysis, we correlate data from all pairs of data types in the same tissue (or cell line) and we correlate all pairs of tissues and cell lines from the same data type. All the results are available from http://stereogene.bioinf.fbb.msu.ru/epiatlas.html. Although this database is large, *StereoGene* compute the correlations efficiently (*≈* 30 sec per each comparison). Therefore, the algorithm well-suited to query inter-sample correlations in large databases. In addition to testing the computational efficiency of *StereoGene* on large databases, the comprehensive analysis enables us to compare the *StereoGene* findings to well-established biological associations.

#### Genome-wide Kernel Correlation analysis

We first use *StereoGene* to assess the correlation between distinct epigenetic tracks from the same tissue type. We focus this part of the analysis on pairwise correlations between the most frequently studied tracks in the Roadmap Epigenomics Project, namely, H3K4me1, H3K4me3, H3K9me3, H3K27me3, H3K36me3 epigenetic features and the RNA-seq and on comparisons of the distributions of the KC values for the same epigenetic tracks across fetal and across adult tissues. The distributions of the correlations are presented separately for fetal tissues and for adult tissues (Figure 2A, Supplementary File 1, Section 10). Generally, the epigenetics marks are more correlated in adult tissues in comparison with the fetal tissues. The statistical significance of this observation is shown on Figure 2,B. The highest difference of feature-to-feature correlation between the collections is observed for the H3K9me3 vs H3K27me3 pair: they are significantly more correlated in adult tissues than in fetal ones. A comparison of correlation between H3K9me3 and H3K27me3 in the same tissue for fetal and adult gave a *p - value* = 3.2 *·* 10*^-^*^5^ (Wilcoxon test). This result is consistent with the prior observation that at early stages, different genomic regions are separately regulated by H3K9me3 and H3K27me3, but during tissue maturation, these heterochromatin marks became more synchronized [5]. One possible explanation is that H3K27me3 initiates chromatin compaction by recruitment of H3K9me3. The colocalization of H3K27me3 vs H3K36me3 relates to the monoallelic gene expression [19]. Fig. 2 also shows a significant increase of correlation of these marks in adult tissues in comparison with fetal tissues. The observation is consistent with the recent studies [20].

**Figure 2.**
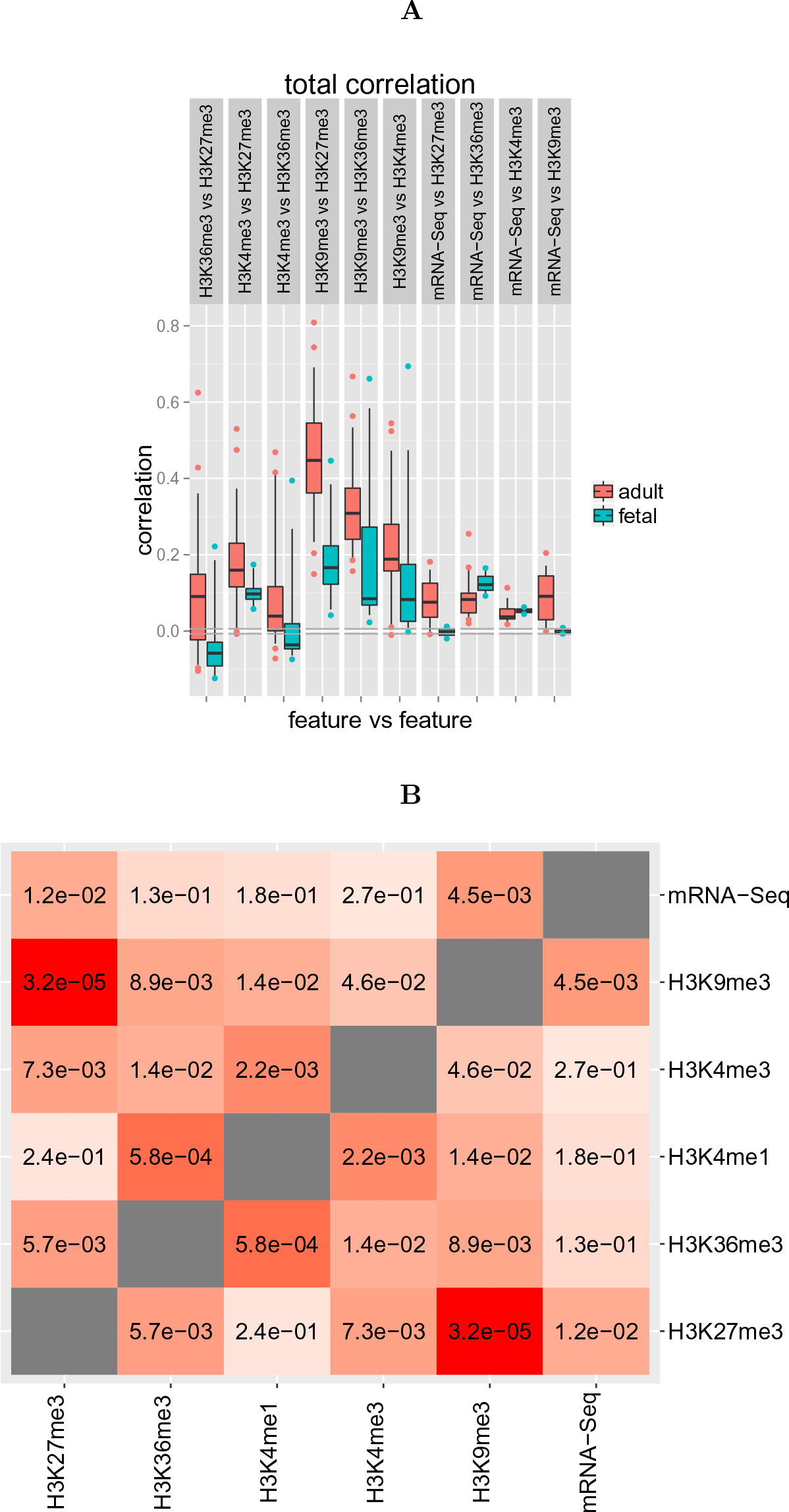
Distributions of genomewide kernel correlations (KC) values of pairs of epigenetic marks across the fetal and across the adult tissues. 12 fetal and 39 adult cell types data are used. A: Boxplots of the KC value distribution for adult and fetal tissues. Gray horizontal lines near the zero show the maximal and the minimal background correlations that are observed over all the datasets. B: p-values for difference of these correlation distributions between fetal and adult tissues (Wilcoxon test)

#### Kernel Correlation windows distribution

To look at the KC difference between adult and fetal tissues in more details, we compare the H3K4me3 and H3K27me3 KC distributions over the windows of the genome in the adult lung tissue, fetal lung tissue and their common background (Fig.3A). In this analysis, we observe a higher correlation between H3Kme4 and H3K27me3 in adult in than fetal tissues. This observation is consistent with chromatin changes during development. Specifically, adult tissues have more regions with “poised promoters” in which both marks are active than do fetal tissues [26].

**Figure 3.**
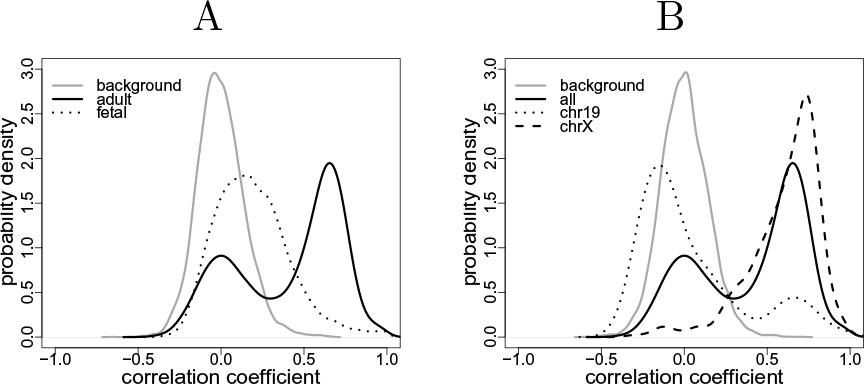
Distributions of correlations. A. H3K27me3 vs H3K4me3 in lung tissues. Solid black line – adult lung; dotted line – fetal lung; gray – the background distribution that coincides for both cell types. B. Correlation distribution for H3K27me3 vs H3K4me3 in female adult lung cells with chromosome specification. Grey – background distribution; solid line – correlation distribution over genome; dashed – correlation distribution for chromosome 19, dotted line – correlations for X-chromosome.

#### Local Correlation analysis

*StereoGene* enables positional interpretation of the correlation resultsproviding the local correlation functionality (see methods). Here, we analyze the local correlation track between the data for H3K4me3 and H3K27me3 marks in adult lung tissue discussed in the previous example. Namely, we compute the local correlation track to define peaks that define the correlation between these two chromatin marks. We then analyze its function through gene set enrichment analysis (as described in the methods) of these peaks in resulting local correlation track (Supplementary file 3). We found that 53 gene sets have FDR ¡5%. In particular, we see the sets “cell motion regulation (FDR ¡ 10*^-^*^4^)” and “positive regulation of cell migration” (FDR ¡ 10*^-^*^3^) that are associated with lung development. Specifically, the cell motion is very active during lung development, then it stops in adult lung, and its regulation genes are poised, i.e. switched off.

#### Cross-correlation analysis. Nucleosome dependency of epigenetic marks

As a part of the standard result, *StereoGene* returns the cross-correlation function between the pairs of samples that were compared. In the Roadmap Epigenomics Project data analysis (see http://stereogene.bioinf.fbb.msu.ru/epiatlas.html), in many cases, we observe the cross-correlation function that has a narrow peak centered at zero. For example, Supplementary Fig. S1 shows the distributions of the KC and the cross-correlation function for tracks H3K27me3 vs H3K36me3 in fetal brain cells. Both H3K27me3 and H3K36me3 are covalent histone modifications, they are positioned inside a nucleosome. The zero-peak could reflect frequent co-occurrence of the marks that results from their co-localization inside one nucleosome. This possibility is supported by recent results from reChIP [13].

#### Simulations. Nucleosome dependency of epigenetic marks

Widespread application of *StereoGene* to epigenetic data indicates that positive correlations between histone marks are far more common than negative correlation (see, e.g., Fig. 2) for all histone marks and tissue types. To test whether the pervasive positive correlations has the same nature as the zero-position peak of the cross-correlation function and that they both occur due to the nucleosome positioning, we perform a simulation experiment (Supplementary File 1, Section 8). We simulate a “genome” that is 60 Mbases long and contains 100000 randomly distributed “nucleosomes”. Then we generate two independent signals that are located only on “nucleosomes”. As a result, we have obtained simulated data with true positive correlation and with zero peaks. When the signals are produced independently of the “nucleosomes”, the peak is not observed and the KC approximately equals to zero. On the other hand, when the simulated signals are co-localized in the simulated nucleosomes we observe a sharp cross-correlation peak at zero. Thus, this simulation suggests that the prevalence of positive kernel correlation values and the zero-positioned peak on the cross-correlation observed above arise from nucleosome positioning rather than from artifacts.

### Chromosome-specific correlation of the histone modifications

The relation between epigenetic marks can differ from chromosome to chromosome. *StereoGene* allows to provide the analysis separately by chromosomes. To show this feature, we compared the relationship between two well-investigated histone marks: the promoter-related H3K4me3, and the heterochromatin polycomb-related H3K27me3, in the adult lung, chromosome by chromosome (see Fig. 3B). The window KC value distribution for these marks over all genome has a high peak on positive correlations. At the same time, the distribution on chromosome 19 has a significantly different shape and it is shifted to lower values. This could be explained by the high gene density, especially, high housekeeping genes density on the chromosome 19. The correlation distribution on chromosome X also differs from the distribution over genome and the chromosome X distribution has a peak on very high correlations (see Fig. 3B), that can be due to the two copies X-chromosome, one is repressed by H3K27me3 mark, while another is active. This observation is consistent with known suppression of chromosome X in female origin somatic cells as part of development.

### Partial correlations

A pair of features in genome never exist in isolation. Interactions of biological features with other genome-wide features will skew the spatial correlation. For example, differences in mappability of short regions to different regions of the genome will impact the signal of all genome-wide data that requires alignment, and the correlations that can be computed in different regions. The influence of additional and sometimes technical genome-wide features sometimes shadows the effect or can cause an apparent new effect that is unrelated to the biology. We call these additional features that impact correlation of biological features “third players” or “confounders”. When the third player genome-wide track is provided, *StereoGene* uses the partial correlation use-case to eliminate its impact on correlation. Here, we provide examples where using this third track enables *StereoGene* to both uncover shadowed effects and remove technical associations between unrelated tracks.

The H3K4me3 is an “active promoter” mark and it is expected to be positively correlated with RNA-seq. Indeed, Fig. 4A shows some weak positive, though statistically significant correlation in Brain Hippocampus Middle. This tissue is an adult one, and we suppose that a significant share of the promoters is “poised” [26] (see the Local Correlation Analysis). In other words, the H3K4me3 effect on the expression is shaded by the H3K27me3 presence. We used the partial correlation mode to remove H3K27me3 influence (Fig. 4B), and the correlation we observe is much stronger. This suggests that the relationship of H3K4me3 to gene expression is modulated by H3K27me3.

**Figure 4.**
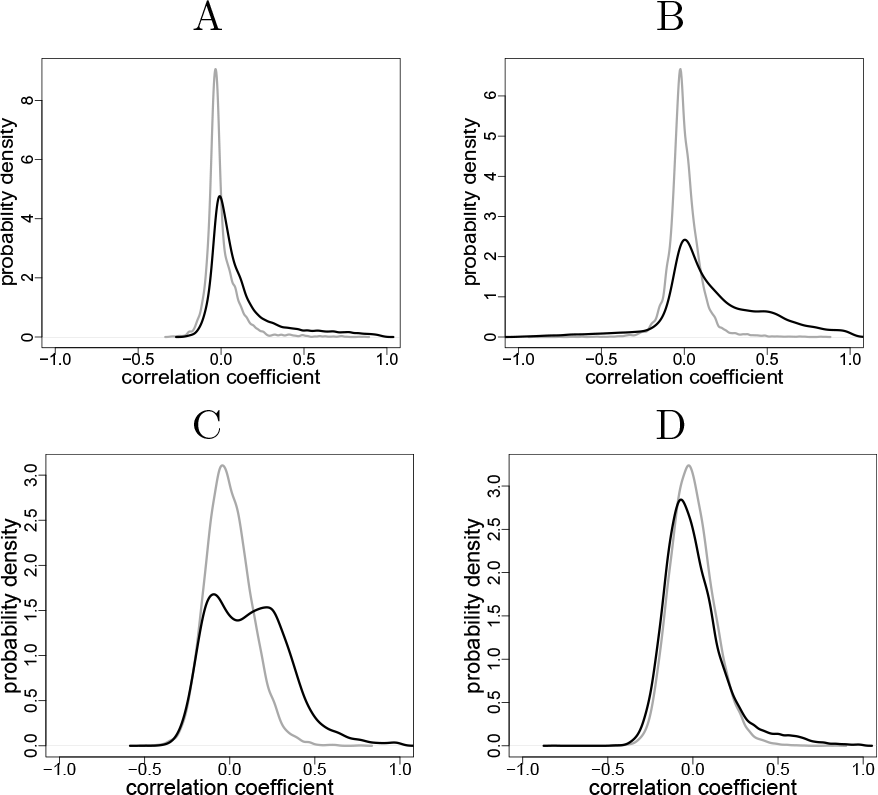
Distributions of the *KC* values over the windows: A: Correlation of H3K4me3 vs mRNA-seq in Brain Hippocampus Middle; B: The same correlation, where the H3K27me3 track is accounted for as a confounder by the partial correlation procedure. C: *KC* values distribution for H3K27me3 vs H3K4me3 in GM12878 cells; D: The same with the nucleosome track accounted for as a confounder. Black line – foreground distribution; gray line – background correlation distribution.

As we observed in the cross-correlation analysis, exact nucleosome positioning confounds all the pairwise histone marks. Therefore, correlations between histone marks often occur because of detection near nucleosome coordinates rather than biological similarities between histone marks in the cell. Using a nucleosome track as the confounder for partial correlation in *StereoGene* decreases the correlation between H3K27me3 and H3K4me3 in GM12878 cells (Fig. 4C-D, Fig. S2). Removing the effect of the nucleosome positioning is limited to datasets that contain nucleosome track data, which are regrettably few. More detailed description about the influence of the confounders and their removal with partial correlation functions in *StereoGene* is presented in the Supplementary File 1, Section 3. Another promising application of the partial correlation to the ChIP-seq data is to exclude the input DNA track as a confounder (Fig. S2)).

### Cross-correlation function: Chromatin marks vs gene features

The local regulation of transcription by chromatin marks usually depends on the positioning of the modified nucleosome relative to the transcription start site. Similar dependence may also occur for other gene features, including gene end sites or exons-introns boundaries. We apply the cross-correlation function in *StereoGene* to assess the relationship of such gene features to expressed and silenced genes in the brain cingulate gyrus. To compute this correlation, we first define a set of expressed genes as those with the top 25% of mRNA-seq gene values of gene counts and silenced genes as those with gene counts in the bottom 25%. All other genes are called moderately expressed. Then, we plot the cross-correlation function of histone marks vs gene features – start/end and intron beginning/end (Supplementary File 1, Section 4) aggregated for each group of genes. We observe the following.

- The distribution of H3K4meX and H3K9ac near TSS of the active genes has two high peaks left and right from TSS and a gap at TSS position. This behavior is in an agreement with other research [8].
- Both H3K4meX and H3K9ac density have a sharp break near intron ends. This behavior may be related to epigenomic splice site definition [4].
- H3K27me3 has a rather narrow peak downstream from TSS of active genes, while for the low-expressed and for the silent genes the peak is wider and it covers TSS. This peak in active genes may be related to a monoallelic expression [19].

### DNA binding proteins: Cohesin and histone modifications

One of the promising applications of *StereoGene* is the analysis of relations of DNA-binding protein with other genomic features. We use the KC from *StereoGene* to determine the positional correlations of cohesin protein Rad21 ChIP-seq track with CTCF track and with different histone modifications in H1 stem cells (RRID:CVCL 9771) and in the K562 (RRID:CVCL 5145) cell line (Supplementary File 1, Section 5). We observe a very strong positional correlation of the CTCF binding with Rad21 binding (p-value *≈* 0, see Supplementary File 1, Section 5, Table 1). Another observation is that promoter and enhancer marks (H3K4meX) are co-localized with cohesin binding (p-value *<* 10*^-^*^16^), while actively transcribed gene regions and repressed gene regions are not. These observations are consistent with [29] and suggest that *StereoGene* is a robust tool to associate DNA protein bindings.

**Table 1.** Types of correlations produced by *StereoGeneS*

| Type of correlation | Eq. | Description |
| --- | --- | --- |
| Kernel correlation (KC) | (3) | The genome-wide correlation coefficient, which is calculated with the kernel; it reflects overall relationship between two tracks. |
| Cross-correlation | (5) | The cross-correlation function shows the structure of the correlation that reflects distance dependence of the tracks values. |
| Local correlation | (6) | This output track shows the Kernel correlation as a function of the genome position. The track can be displayed in genome browsers and it can be used in further analysis. |
| Partial correlation | (7) | The correlation computed between a pair of profiles, excluding the impact of a third confounding genomic track. |

### Genomewide expression: CAGE vs gene annotation

CAGE FANTOM4 [24] data (CAGE clusters) represents a genomewide map of capped mRNA. The CAGE data is expected to estimate mRNA that are prevented from degradation and promoted for translation genomewide. As a result, these data are hypothesized to correlate strongly with transcription start sites. To determine whether there is a statistically significant CAGE signal in gene start sites and other gene features, we analyze the positional relationship of CAGE data, for the nucleus and for cytosol of H1-hESC cells with the RefSeq [22] gene annotations with the *StereoGene* Kernel correlation. As hypothesized, CAGE clusters are highly correlated with gene starts. We do not see any signal at the promoter regions, but we observe less obvious phenomena, namely, strong positional correlation of CAGE clusters with the intron start sites (donor splice sites) and strong positional correlation of CAGE clusters with transcription termination sites, see Supplementary File 1, Section 5 for more details. CAGE association with intron starts may be explained by the activity of debranching enzymes [25]. After lariat debranching, the freed 5′ end of the intron may become available for capping, and this cap would be detected by CAGE. Indeed, short (18-30 nucleotides) RNAs with the 3’-end that exactly maps to donor splice sites are observed [30]. The transcriptional termination site correlation is less evident, though it suggests that occasional capping of the free 5′ end after cleavage by the polyadenylation complex is possible. The *StereoGene* analysis of CAGE data enables unprecedented associations of CAGE with gene annotations to assess the function on mRNA capping in different gene features.

### Comparison with other methods

We compare (Table 2) the *StereoGene* functionality with that of commonly used tools. Notably, very few programs can compute on continuous data and require the establishment of often arbitrary thresholds in order to create intervals for analysis. KLTepigenome [18] is able to work with the continuous profiles, but it is limited to sparse data and is quite slow even when being compared to *StereoGene* doing the same computation on the full profile.

**Table 2.**
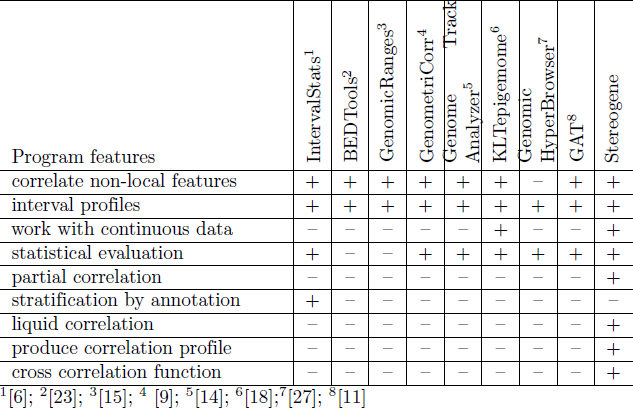
Comparison of functionality for correlation analysis programs.

We test consistency of our results with the results, which are described in [33] on modENCODE [10] S2-DRSC cell line dataset. We find the numeric agreement to be satisfactory. The Pearson correlation coefficient of the Kernel Correlation values and the Interaction energy score [33] is 0.48. Our results are summarized and visualized in the Supplementary File 1, Section 7.

## Discussion

We present a new method, *StereoGene*, with unprecedented speed for estimation of genomewide positional correlations. A comprehensive description of the mutual positioning of genomewide tracks requires a set of statistical test on different scales. For that to happen, *StereoGene* provides a collection of genome-wide correlation techniques (see Results).

We apply the program for a variety of datasets including continuous (ChIP-seq) and interval (genome annotation) data. The results are consistent with recent biology knowledge. In addition, we observe the changes in the correlation between epigenomic features across developmental trajectories of several tissue types, and we find an unexpected strong spatial correlation of CAGE clusters with splicing donor and poly(A) sites. Both observations require verification and both are in concordance with the biological intuition of other authors.

In contrast with other methods in the literature, *StereoGene* is unique in its ability to rapidly compute correlations of continuous genome-wide features in addition to discrete gene intervals used in most correlation techniques. As seen on public datasets, the approach yields biologically plausible results. The most common application of *StereoGene* is the association of distinct genomic features from the same individual or common genomic feature between individuals. The correlation distribution plots enable assessment of directionality in addition to statistical significance, to depict multiple varieties in these genomewide associations. In addition, local correlation tracks can be used for traditional gene enrichment analysis or to describe the relationship between genomic features. The partial correlation allows excluding of a known confounder. Other features of *StereoGene*, such as batch analysis and using of linear models, make this tool useful for mass and diverse analysis of the genomic tracks.

As far as we work with the continuous tracks directly, we do not lose information on the binarization approaches as the most methods do. In these other methods, the choice of the binarization threshold is usually supported by reasonable statistical considerations. However, despite the threshold quality, the dependency of the results the threshold should be checked before conclusions of biological associations are drawn with these threshold-based methods. The biological data are obtained on a large population of cells, which can be very inhomogeneous, see, e.g. the phenomenon of gene expression bursts [2]. Thus, even small averaged signals can be biologically significant. We test (Supplementary File 1, Section 9) the dependency of the correlation (KC) value on the binarization threshold. High thresholds lead to overestimated correlations.

Currently, *StereoGene* is widely applicable for analysis of similarity of genomic-track-represented biological data, including massive analysis. The track-to-track distance results can be aggregated to compare different tissues and different time course points.

Quite often, we observe bimodal KC distribution, and a question whether the modes correspond to some global chromatin states is naturally raised. The first hypothesis we intend to test in this way is a relation to the chromatin A/B compartmentalization [7]. The *LC* track can be compared with some third data source by the next run of *StereoGene*; the result of the sequential runs is a 3-way correlation that is analogous to the liquid correlation [16]. This analysis will enable, particularly, more fine testing of the relations of epigenetic features mutual positioning along the chromosome with the 3D positioning of the chromatin.

We intend to add statistical tests that compare the distributions of observed correlations from different track pairs (e.g. input tracks). It is a natural differential mode without the permutation-based estimations. For the case when the researcher has a collection of tracks for the same tissue, we plan to computationally estimate their common component to use it as a common input track. These new approaches will extend *StereoGene* beyond robust genome-wide associations to a comprehensive platform for continuous data analysis of genome-wide tracks for cross-platform, integrated genomics analyses.

## Acknowledgments

We are grateful to Roman Kudrin, Ekaterina Khrameeva and Alexandra Galytsyna for testing the program. Thanks to Renat Arufilov, Artur Zalevsky and to Dmitriy Vinogradov for technical solutions and for support. Thanks to Aleksey Stupnikov for his ideas for the future. Thanks to Leslie Cope for his advice. Thanks to Patricia Palmer for her help with the text of the manuscript.

### Funding

This work was supported by Russian Science Foundation (grant 14-24-00155), by National Institutes of Health (grants P30 CA006973 and NCI R01CA177669), by Allegheny Health Network-Johns Hopkins Cancer Research Fund and JHU IDIES/Moore Foundation and by Russian Foundation for Basic Research (grants 14-04-01872 and 14-04-00576)

